# Both composition and configuration of forests and urban development shape bat activity and diversity in North American temperate forests

**DOI:** 10.1101/2025.02.14.638160

**Authors:** Sihao Chen, Han Li

**Affiliations:** Department of Health and Environmental Sciences, Xi’an Jiaotong-Liverpool University, Suzhou 215123, China; Department of Earth, Ocean and Ecological Sciences, School of Environmental Sciences, University of Liverpool, Liverpool L69 3BX, United Kingdom; Department of Biology, University of Nebraska Omaha, 6220 Maverick Plaza AH114, Omaha, 68182, Nebraska, United States

**Keywords:** Forests, Bats, Chiroptera, Urban, Landscape index, Composition, Configuration

## Abstract

Temperate forest ecosystems are important habitats for many bat species. However, these habitats are increasingly affected by anthropogenic disturbances, particularly urban development, leading to landscapes with varying land cover composition and configuration. Limited research has examined how forest and urban landscape composition and configuration influence bat activity and diversity. Using a multi-year statewide bat acoustic monitoring dataset from North Carolina, United States, we investigated the effects of forest and urban composition and configuration at multiple spatial scales on bat activity and diversity. First, we constructed single-variable landscape index regression models and found that both composition and configuration of forests and urban developments influenced bat activity and diversity in a species-specific manner. Next, we applied a hierarchical partitioning approach to compare the relative contributions of composition and configuration indices in explaining variance in bat activity. For big brown bats and hoary bats, evergreen forest and urban development composition indices contributed the most to explaining activity variance. In contrast, for eastern red bats, evening bats, and tricolored bats, deciduous forest fragmentation indices describing landscape configuration were the most influential factors. Silver-haired bat activity variance was primarily explained by an evergreen forest fragmentation index. Lastly, urban development configuration indices were the strongest predictors of Mexican free-tailed bat activity and total bat activity. These results suggest that forest and urban landscape configuration should be considered in conservation and management planning for North American temperate forest ecosystems, particularly in regions that have not experienced drastic deforestation in recent decades.

## 1. Introduction

Forests provide essential ecosystem services such as timber production, carbon sequestration, recreational opportunities, and biogeochemical cycling, while also harboring rich biodiversity (Brockerhoff et al. 2017). However, increasing evidence suggests that forests are experiencing significant disturbances due to human activities, particularly habitat modification for agriculture and urban development (Haddad et al. 2015; Li et al. 2022). Consequently, forest habitats become highly fragmented, resulting in forest remnants with varying configurations. This fragmentation threatens the ability of the forest ecosystems to maintain their composition (i.e., abundance of different forest types), configuration (i.e., the spatial pattern and characteristics of different forest patches), and overall species biodiversity (Haddad et al. 2015; Newbold et al. 2015). Many species, such as bats that reply on forests for food and shelter, are heavily affected because altered forest composition and configuration across various spatial scales can lead to unsuitable habitats (Klingbeil and Willig 2009; Russo and Ancillotto 2015; Frick et al. 2020). Changes in habitat suitability, coupled with increasing urbanization, make it challenging to develop comprehensive forest management strategies that effectively preserve forest habitats and conserve species at both site and landscape scales (Pfeifer et al. 2017; Arroyo-Rodríguez et al. 2020). Therefore, understanding how species respond to altered landscape composition and configuration across spatial scales is essential for forest biodiversity conservation (Haddad et al. 2015; Newbold et al. 2015).

Bats are the second most diverse mammal order, with over 1400 species worldwide (Frick et al. 2020). Bats provide essential ecosystem services such as pest control, pollination, and biogeochemical cycling (Kunz et al. 2011). However, habitat loss and degradation are major threats to bats and must be addressed for effective bat conservation (Frick et al. 2020). Forests serve as critical roosting and foraging habitats for many bat species (Lacki et al. 2007). The loss of forest habitats alters land cover composition in the landscape, contributing to bat population declines in forest ecosystems (Arroyo-Rodríguez et al. 2016; Frick et al. 2020). Reductions in deciduous, evergreen, or mixed forest areas directly impact bat occurrence and activity by affecting foraging and roosting habitat availability (Kalcounis et al. 1999; Lacki et al. 2007). For example, deciduous forests containing grouped tree types such as oak-hickory (*Quercus* spp., *Carya* spp.) or maple-beech-birch (*Acer* spp., *Fagus* spp., *Betula* spp.) are different from evergreen forests, such as spruce-fir (*Picea* spp., *Abies* spp.) and pine (*Pinus* spp.), in terms of understory vegetation, deadwood availability, and clutter amount due to seasonal leave shedding (McElhinny et al. 2005; Brown and Lambert 2018). As a result, deciduous forests support rich understory invertebrate communities (e.g., beetles, flies, and sawflies) and provide diverse snags, logs, and litter conditions for foraging and roosting (Rudolph et al. 2009). Beyond forest habitat loss, the expansion of urban areas within forest ecosystems creates new ecological niches that favor generalist species over forest specialists (Russo and Ancillotto 2015; Li and Kalcounis Rueppell 2018). For example, in Panama, *Molossus spp.*, *Cynomops spp.*, and *Eumops spp.* were detected more frequently in urban areas than in adjacent forest habitats (Jung and Kalko 2011). Recent studies also indicate that higher building density and specific anthropogenic structures might facilitate the occurrence of certain bat species such as Mexican free-tailed *Tadarida brasiliensis* in urban areas, expanding the distribution range (Li and Wilkins 2014, 2015; McCracken et al. 2018).

Forest habitat loss also leads to habitat fragmentation, resulted in a higher number of relatively small patches, reduced habitat connectivity, and altered landscape configuration across spatial scales (Ethier and Fahrig 2011; Pfeifer et al. 2017). This process can further create a range of habitats with diverse edge conditions (Fahrig 2017). Changes in forest configuration have mixed effects on bats. At the urban-forest interface, higher foraging activity has been documented compared to either urban or forest habitats, suggesting that edge habitats might serve as more preferable foraging grounds for Neotropical aerial insectivorous bats (Jung and Kalko 2011). Additionally, urban fringes and agricultural lands near urban areas provide valuable complementary resources (e.g., diverse insect prey) for highly mobile bat species that have adapted to roost in human-made structures such as bridges, tunnels, and attics, where thermal stability and predator protection are advantageous (Li and Wilkins 2015; Aguiar et al. 2021; Li and White 2024). On the other hand, forest fragmentation can negatively impact forest-dependent species (Jung and Kalko 2010; Threlfall et al. 2011; Russo and Ancillotto 2015). Connectivity between habitat patches is particularly critical for certain bat species in fragmented and complex landscapes. Additional forest corridors and linear infrastructure, such as greenways and transportation corridors in urban areas, can enhance habitat suitability and support species that require well-connected landscapes for survival and mitigate the negative effects of isolation (Straka et al. 2019; Carlier et al. 2019). These linear features are often associated with hedgerows and early to mid-successional habitats, which facilitate commuting between foraging and roosting sites in complex habitat mosaics (Tournant et al. 2013; Straka et al. 2019; Cable et al. 2021).

Landscape habitat composition and configuration changes are interconnected processes (Haddad et al. 2015; Arroyo-Rodríguez et al. 2016; Fahrig 2017). Limited research has comparatively examined their effects on bats. Existing literature highlights the importance of both habitat composition and configuration, as well as species- or guild-specific responses in bats. For example, Klingbeil and Willig (2009) found that in the Amazon Basin, frugivorous bats were more responsive to landscape composition due to variations in food availability, whereas gleaning animalivores were more influenced by landscape configuration due to movement restrictions. In the United Kingdom, broadleaved woodlands were important for bat roost locations in terms of both extent and proximity (Boughey et al. 2011). In Chiapas, Mexico, Arroyo-Rodríguez et al. (2016) found that the forest cover amount was more influential than the degree of forest fragmentation in explaining phyllostomid bat assemblages. Geographically, the effects of forest habitat composition and configuration on Neotropical bats have been more extensively studied (Klingbeil and Willig 2009; Rocha et al. 2016; Arroyo-Rodríguez et al. 2016; Falcão et al. 2021). Similar to tropic forests, temperate forests, an essential component of global forest ecosystems, are also highly susceptible to human-induced landscape changes (Newbold et al. 2015; Li et al. 2022). Bats in temperate forests are also experiencing population declines due to forest habitat loss and fragmentation (Lacki et al. 2007; Law and Blakey 2021). However, no research has specifically investigated how changes in forest habitat composition and configuration affect North American temperate forest bats or whether landscape composition or configuration comparatively has a stronger influence on bat populations. This gap in knowledge limits the development of more effective forest conservation and management strategies.

When examining the effects of forest habitat loss and fragmentation, it is essential to consider varying species responses to the same environmental factor across spatial scales. A well-established effect at one spatial scale may not necessarily be extrapolated to other scales (Turner et al. 2001; Fletcher Jr. et al. 2023; Fahrig 2024). For example, bat activity of forest foragers from Family Vespertilionidae and Emballonuridae was positively associated with more forest cover (mature or secondary) at a 500 m scale in northeastern Brazil; however, this positive effect was not observed at a 2000 m scale (Falcão et al. 2021). Conversely, Conversely, in central Chile, a study found that the positive relationship between forest cover amount and *Myotis chiloensis* at a 1000 m scale became negative at a 4000 m scale (Rodríguez-San Pedro and Simonetti 2015). Regarding the effects of forest configuration, a higher number of forest patches led to increased activity of *Myotis crypticus, Lasiurus cinereus, Myotis chiloensis,* and *Tadarida brasiliensis* at multiple scales (500 m to 2500 m), while it decreased the activity of *Pipistrellus pipistrellus* at the 500 m scale (Rodríguez-San Pedro and Simonetti 2015; Laforge et al. 2022). Similar variations across spatial scales have also been observed in bats inhabiting urban areas (Gallo et al. 2018; Moretto et al. 2019). Therefore, a comprehensive assessment of the complex effects of landscape composition and configuration across multiple spatial scales is essential to understanding species responses in temperate forest ecosystems. This understanding is crucial for shaping effective forest management policies and decisions (Fletcher Jr. et al. 2023).

In this research, we used a multi-year statewide bat acoustic monitoring dataset from North Carolina, United States, to investigate how the composition and configuration of urban and forest lands at multiple spatial scales influenced bat activity and diversity in the temperate forest ecosystem. Our first objective was to determine which landscape indices of urban and forest composition and configuration affected total bat activity, species richness, and species-specific activity, as well as how they exerted these effects. Our overarching hypothesis for this objective was that different species would respond to specific landscape indices in ways that reflected their functional traits and ecological needs. Specifically, we hypothesized that: 1) Tree-roosting bats would positively correlate with forest composition indices, whereas bats that roosted in anthropogenic structures would positively correlate with urban composition indices (Arroyo-Rodríguez et al. 2016; Li and Kalcounis Rueppell 2018); 2) forest configuration indices would positively correlate with some tree-roosting bat species (Ethier and Fahrig 2011), whereas urban configuration indices might not affect bats, as no research has reported such patterns; 3) due to species-specific responses to urban and forests, no correlative patterns would be found for total bat activity and species richness as they described all bats collectively (Klingbeil and Willig 2009; Boughey et al. 2011). Additionally, we aimed to examine whether the effects of landscape indices were consistent across spatial scales under this objective. Our second objective was to compare the relative influence of landscape composition and configuration on total bat activity, species richness, and species-specific activity. To achieve the objective, we employed a hierarchical partitioning approach. First, we compared the relative importance of composition and configuration indices within specific land cover types (deciduous, evergreen, mixed forest, and urban development). Next, we compared landscape indices across different land cover types to identify the most influential factor shaping total bat activity, species richness, and species-specific activity in temperate forest ecosystems. Given the conflicting findings in the existing literature (Ethier and Fahrig 2011; Arroyo-Rodríguez et al. 2016), we hypothesized that species-specific patterns would emerge regarding whether landscape composition or configuration had a greater influence on bat activity.

## 2. Methods

### 2.1. Study area

The research area was within the state of North Carolina, United States. North Carolina belongs to the eastern temperate forests ecoregion with three distinct sub-ecoregions (Omernik and Griffith 2014), commonly named as mountain, piedmont, and coastal plain. According to the United States Forest Service, approximately 60% of the land area of North Carolina is forest land with a relatively stable trend in the total forest area (Brown and Lambert 2018). The most common forest-type group is oak-hickory, followed by loblolly-shortleaf, oak-pine and oak-gum-cypress, *Nyssa* spp., *Taxodium* spp., all of which can be found in three regions within North Carolina. The majority (>90%) of forests in North Carolina are timberland, mostly (>80%) owned privately by individuals or nonindustrial entities (Brown and Lambert 2018). In the past two decades, North Carolina has been experiencing rapid human population growth (U.S. Census Bureau 2021) and related urbanization (Yang et al. 2018). The piedmont region has the largest cities, even though increased urban developments are seen throughout the state. North Carolina has a total of 17 bat species, among which at least eight species have a statewide distribution (Li and Kalcounis Rueppell 2018; Li et al. 2024). North Carolina has a long history of baseline bat monitoring, and since 2015, North Carolina has been implementing the North American Bat Monitoring Program (NABat) statewide (Li and Kalcounis Rueppell 2018; Reichert et al. 2021).

### 2.2. Bat acoustic monitoring data collection and processing

We used publicly available NABat mobile transect survey data from 2016, 2019, and 2021 in North Carolina. The NABat mobile transect survey occurs annually between June and July. This survey effort follows the NABat grid cell masterplan and site selection guide to choose survey sites (Loeb et al. 2015; Reichert et al. 2021). A site in our research is a 10km by 10km square cell (named as a NABat grid cell). Within a cell, a 32km transect on public roads was mapped to maximize its coverage for all possible land cover types along the road within a cell. Detailed transect design was described in previous studies (Li and Kalcounis Rueppell 2018; Li et al. 2019). In June and July each year, a transect is surveyed twice within 7 nights by either professional wildlife technicians or community science volunteers. All surveyors were trained prior to the field season on equipment usage and survey protocols. The survey time window for each transect in June or July is kept relatively consistent across years (e.g., a transect was surveyed in early June in 2016, it would be surveyed again in early June in both 2019 and 2021).

On a sampling night, the transect survey started 45 minutes after sunset. A vehicle traveled at a 32km/hr speed to cover the transect, following local traffic laws. A bat detector was attached to the vehicle roof to record bats passing over the vehicle. In 2016, AnaBat SD2 (Titley Scientific Inc., Australia) was used in all surveys. In 2019 and 2021, Echometer Touch (Wildlife Acoustics Inc., United States) was used with a tablet (either Android or IOS based). The AnaBat detector setting was described by Li et al. (2019). The Echometer Touch setting was the manufacturer’s default setting. The detector difference was considered and included as a survey covariate in the statistical analysis. Additional survey covariates included temperature, windspeed, humidity, cloud cover, collected on each survey night via weather information apps on mobile devices. All bat acoustic recordings were stored on external hard drives for processing.

To process acoustic recording files, we first converted 2019 and 2021 acoustic files from full spectrum files to zero-crossing files by AnaBat Insight (Titley Scientific Inc., Australia) to be consistent with 2016 data. We defined a bat pass as a recording sound file with at least three clear and completed calls less than 0.5s apart. Next, we manually removed all sound files that did not qualify as a bat pass. Using all bat passes, we calculated one dependent variable total bat activity (number of passes per transect per night). Following the manual identification protocol described by Li and Kalcounis Rueppell (2018) and Li et al. (2019), the last author identified bat passes to species in 2024. We considered all 17 species possibly present in North Carolina for manual identification except for the Seminole bat (*Lasiurus seminolus*). The Seminole bat cannot be distinguished acoustically from the eastern red bat (*Lasiurus borealis*) in this region, based on existing literature (e.g., Loeb et al. 2008) and the last author’s field experience. We treated these two species as a group, even though we referred to the group as the eastern red bat throughout the document due to the rare records of the Seminole bat in North Carolina.

Based on the manual identification results, we calculated the following dependent variables: species-specific bat activity (number of passes per species per transect per night) and species richness (total number of species per transect per night). For the species-specific bat activity, we included the following species for the statistical analysis (four-letter species abbreviations used in figures): big brown bats (*Eptesicus fuscus*, EPFU), eastern red bats (*Lasiurus borealis*, LABO), hoary bats (*Lasiurus cinereus*, LACI), silver-haired bats (*Lasionycteris noctivagans*, LANO), little brown bat (*Myotis lucifugus*, MYLU), evening bats (*Nycticeius humeralis*, NYHU), tricolored bats (*Perimyotis subflavus*, PESU), and Mexican free-tailed bats (*Tadarida brasiliensis*, TABR). For the species richness, we counted all species that were identified on a transect on each night. As all transects were similar in length by design, we did not further use transect length to standardize dependent variables.

### 2.3. Landscape indices

To estimate the landscape composition and configuration indices, we generated polygon buffers along each transect. We used areas within the circular buffers with radii of 500, 1000, 2500 and 5000 m around transects to extract land cover data. These buffers distances represent the typical home ranges and foraging distances of bats in temperate forests (Ethier and Fahrig 2011; Langridge et al. 2019; Laforge et al. 2022) and were selected to account for the varying responses of different bat species to different landscape scales. In addition, these buffer distances were commonly used in bat studies on the effects of landscape composition and configuration and allow for broad comparison in different landscape types (Ethier and Fahrig 2011; Rodríguez-San Pedro and Simonetti 2015; Rocha et al. 2016; Falcão et al. 2021; Laforge et al. 2022; López-Baucells et al. 2022). Buffer radii greater than 5000 m were not included to minimize extent overlapping and spatial autocorrelation.

We investigated four land cover types: deciduous forest, evergreen forest, mixed forest, and urban development. We used the 30-m spatial resolution National Land Cover Databases in 2016, 2019 and 2021 to quantify landscape composition and configuration indices (Yang et al. 2018). For the National Land Cover Data, we combined all urban development land cover types and collectively considered it as urban. The landscape indices encompassed four broad categories, area, edge, connectivity, and degree of fragmentation (McGarigal et al. 2023). We used area indices to describe landscape composition and the other three categories for landscape configuration, because these indices were widely recognized for their associations with the effects of landscape composition and configuration on biodiversity (Fahrig 2017; Brändel et al. 2020). Specifically, in each polygon buffer, following protocols described by (McGarigal et al. 2023) we calculated three **area-related** indices: mean patch size of a land cover type (AREAMN), total area of a land cover type (CA), and percentage of a land cover type (PLAND); two **edge-related** indices: mean of perimeter-area ratio for a land cover type (PARAMN) and area-weighted mean of shape index (SHAPEAM, a straightforward measure of shape complexity, McGarigal et al. 2023); three **connectivity-related** indices: aggregation index (AI, the frequency of like adjacencies between the same patch land cover type appearing side-by-side on the map), patch cohesion index (COHESION, a measure of the physical connectedness of the corresponding land cover type), and mean contiguity index of a land cover type (CONTIGMN); and **three fragmentation-degree-related** indices: largest patch index (LPI, a dominance measure of the percentage of total landscape area comprised by the largest patch of a land cover), number of patches of a land cover type (NP), and patch density of a land cover type (PD). The indices were calculated with ArcGIS 10.8, arcpy in Python 2.7 and Fragstats v4.2 (McGarigal et al. 2023).

### 2.4 statistical analysis

All statistical analyses including data manipulation and visualization were conducted in R (version 4.2.2, R Core Team 2022). We used p < 0.05 as the significance criterion for all analyses where a p value was generated. Prior to formal analyses, we conducted a series of preliminary analyses to generate base models to include appropriate covariates. Generally, we followed the model construction process described by Li et al. (2019) for the preliminary analysis to generate a base model for each dependent variable. A base model included all necessary covariates and was tested to account for the most suitable dependent variable distribution and autocorrelations. Landscape index independent variables were added in the base model for the formal analysis described below. All numeric independent variables and survey covariates were standardized to the same scale using R package “mosaic”, allowing comparable regression coefficients (Gelman and Hill 2006; Pruim et al. 2017). For dependent variables, we used the variance-to-mean ratio to determine variable distributions. We found that total bat activity and all species-specific bat activities had a variance-to-mean ratio larger than 4. Thus, these dependent variables were modeled with a negative binomial distribution (Gelman and Hill 2006). Species richness had a variance to mean ratio smaller than 4 and was modeled with a Poisson distribution.

To generate a base model, we first tested for temporal autocorrelation in each dependent variable between two nights within a year as described by Li et al. (2019). Consistent with previous work, no temporal autocorrelation was found between nights at a site within a year. We next tested the effect of year, treating it as a categorical variable (2016, 2019, 2021). This was to account for both biological differences due to annual bat migration/habitat change and the bat detector brand difference. We found the effect of year in all dependent variables except for species richness and hoary bat activity. For base models where the effect of year was significant, we constructed generalized linear mixed models (GLMM) in which year was a random effect. For species richness and hoary bat activity, we constructed generalized linear models (GLM). We then tested survey covariates regarding weather and only included significant survey covariates in each base models. Lastly, we tested potential spatial autocorrelation via Moran’s I tests using R package “ape” (Paradis et al. 2004). We detected spatial autocorrelation in total bat activity and species-specific bat activity for the eastern red, evening, and tricolored bats. We followed the method described by Li et al. (2019) to calculate a spatial covariate to include in the base model accordingly to account for spatial autocorrelation. Additionally, we examined if landscape index variables might be correlated with covariates and did not find such correlations.

Once we constructed the base model for each dependent variable, we added one landscape index independent variable at a time to form single variable landscape index models. For a dependent variable, there were four land cover types (deciduous forest, evergreen forest, mixed forest, and urban), 11 landscape indices per land cover type, and four spatial scales per landscape index (176 single variable models per dependent variable). For each single landscape variable model, we extracted the regression coefficient for the landscape index, the corresponding p value, as well as model AIC (model results available in Supplementary material 1). To collectively visualize these model results, we generated a tile graph, in which we color-noted models where the landscape index was significant. We used blue to indicate positive correlations and red for negative correlations, and a dot whose size indicated regression coefficient’s absolute value. As all landscape index independent variables and covariates were standardized as mentioned above, the regression coefficients visualized in the tile graph were directly comparable.

To further understand how forest and urban landscape composition and configuration affected bat activity and diversity comparatively, we constructed multivariate models next. As landscape indices were generally intercorrelated across spatial scales and among land cover types, we detected multicollinearity in our landscape index independent variables. Therefore, we applied a hierarchical partitioning approach to multivariate models to compare relative contribution of independent variables in explaining the variance in dependent variables via partitioning goodness-of-fit for all potential models within a multivariate regression (Smith et al. 2009; Lai et al. 2022, 2023). To ensure each multivariate model was not overwhelmed by too many independent variables (Gelman and Hill 2006), we first constructed models to compare and select composition and configuration indices within a land cover type. For a forest type or urban development, we used single variable model results to choose which landscape index variable to include in the final multivariate model. Using the same base model for a dependent variable, we included all indices that were found significant in the single variable landscape index models. When a landscape index was found significant across multiple scales, we chose the spatial scale that had the largest absolute value of regression coefficient. Through the within-land cover type hierarchical partitioning, we selected the landscape index with the highest contribution to the overall marginal R^2^ for the cross-land cover comparison. For each cross-land cover multivariate model, there would be one landscape index from each forest type and urban development as well as any covariates chosen for the base model. Hierarchical partitioning was conducted with R package “glmm.hp” and its extension based on the distribution of the dependent variable (Lai et al. 2022, 2023). Within-land cover type hierarchical partitioning results were reported in Table 1 and Supplementary Material 1. Cross-land cover type hierarchical partitioning results were visualized as the final model R^2^ and percentage of individual contributions of landscape indices and covariates. Additionally, for landscape indices that contributed the most to R^2^ in each final cross-land cover multivariate models, we visualized the correlation relationship between the landscape index and the corresponding bat dependent variable by plotting the model predicted correlation trend and raw data points.

**Table 1.**
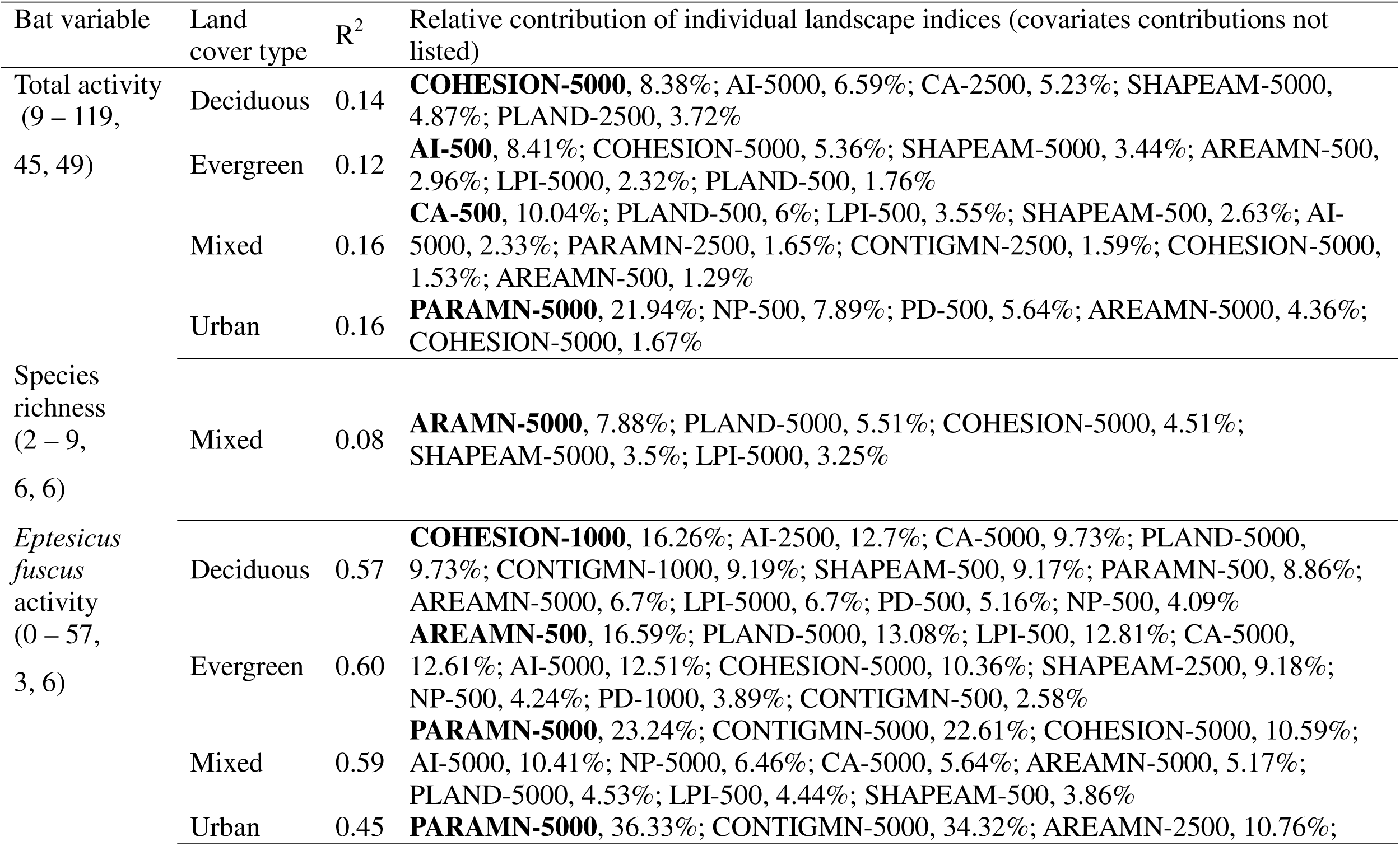
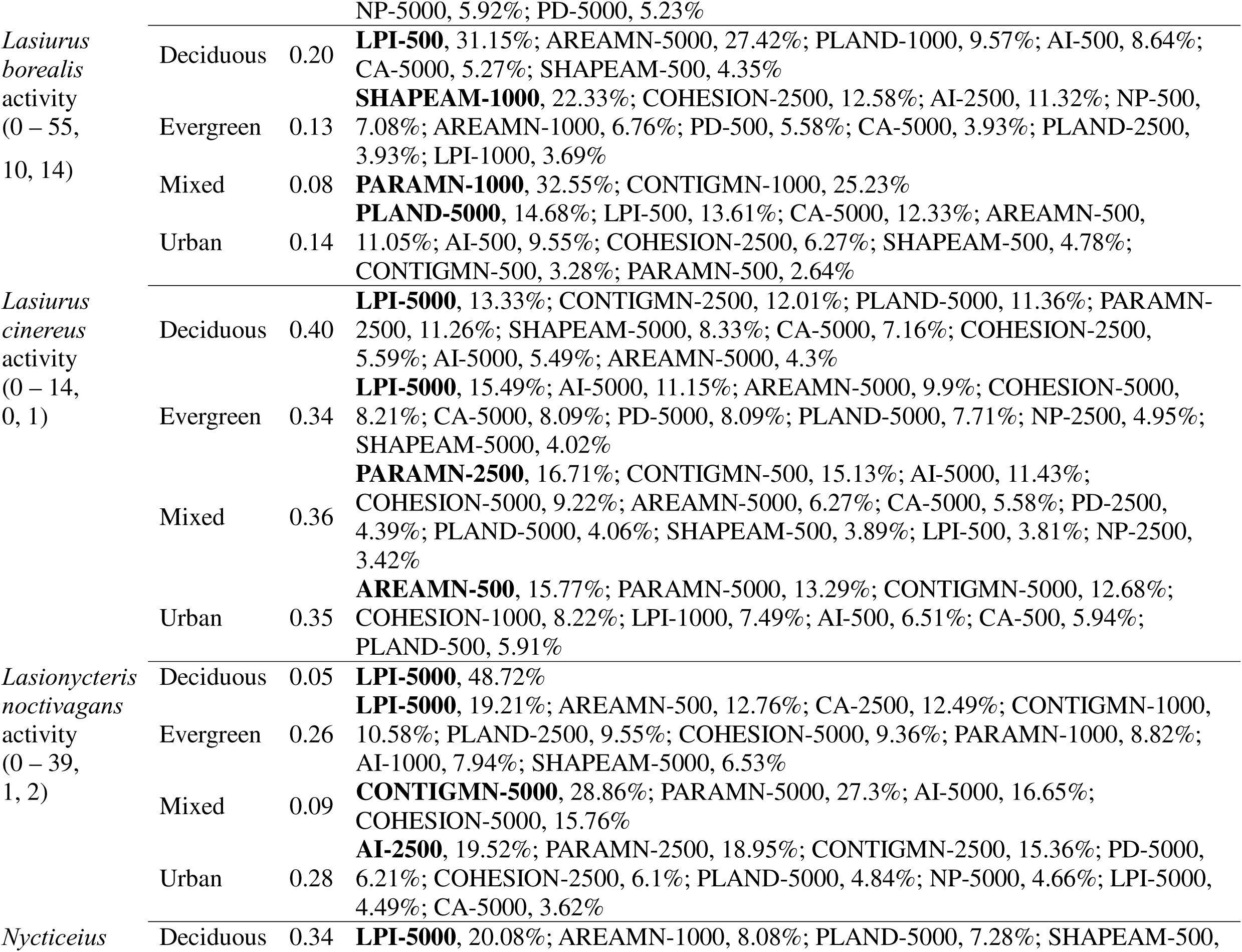
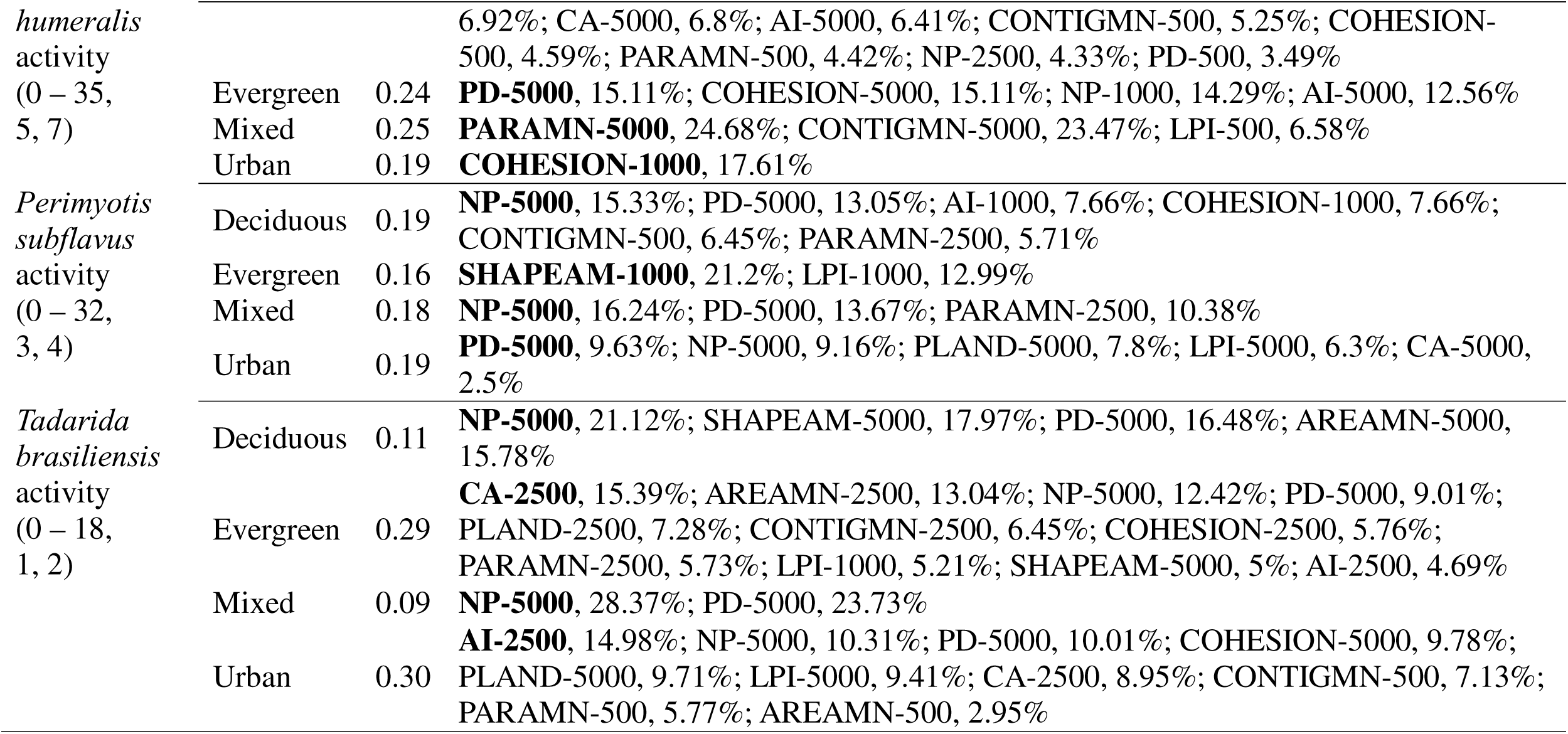
Range, median, and mean of bat dependent variables investigated and within-land cover multivariate model hierarchical partitioning results for relative contributions of landscape indices (spatial scale in number following the dash line, top contributing landscape index in bold).

## 3. Results

The NABat baseline monitoring generated a total of 270 nights of surveys from 54 unique grid cells/transects (Fig. 1). Respectively, 42, 43, and 50 grid cells yielded two nights of survey in 2016, 2019, and 2021, with 32 grid cells that yielded data for all three years. A total of 20,768 recording files were collected from the surveys, among which 13,257 files were considered as individual bat passes and 10,289 passes were identified to 13 species. The species richness on a transect on a night ranged from two to nine species, with a median and a mean of six species. The most commonly recorded bat species was the eastern red bat (3,820 passes), followed by the evening bat (1,802 passes), the big brown bat (16,60 passes), the tricolored bat (1,211 passes), the silver-haired bat (536 passes), the Mexican free-tailed bat (496 passes), and the hoary bat (301 passes). The other species recorded but not included in the quantitative analysis (except for counting in species richness) were the Rafinesque’s big-eared bat (*Corynorhinus rafinesquii*), the northern yellow bat (*Lasiurus intermedius*), the southeastern myotis (*Myotis austroriparius*), the gray bat (*Myotis grisescens*), the little brown bat (*Myotis lucifugus*), and the eastern small-footed myotis (*Myotis leibii*). The total bat activity ranged from nine to 119 passes per transect per night, with a median of 45 passes per transect per night and a mean of 49 passes per transect per night. Species-specific activity range, median, and mean were reported in Table 1.

**Fig. 1.**
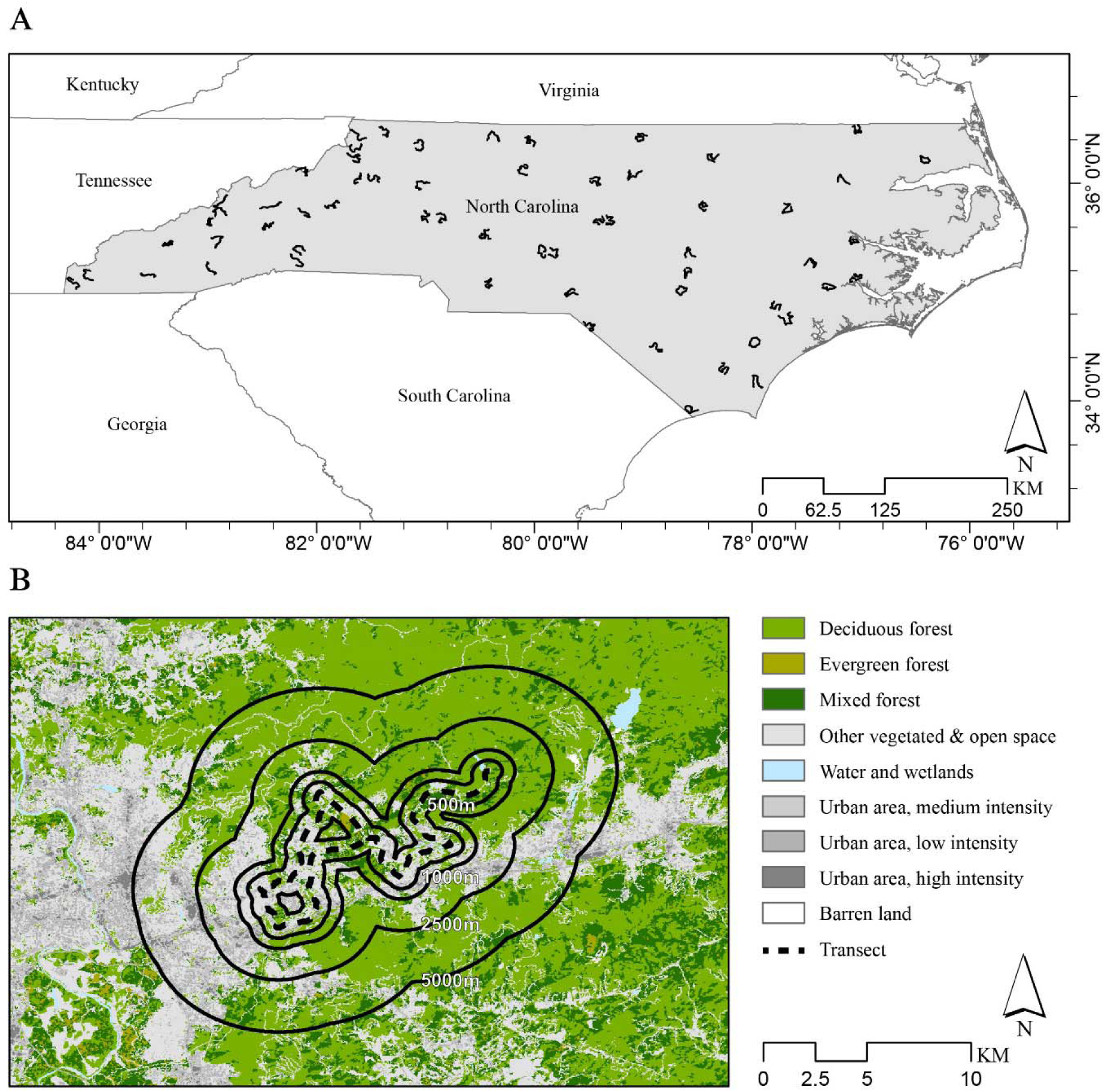
Research area map for the state of North Carolina, United States with the North American Bat Monitoring Program (NABat) transects in blackline (A) and a detailed example of a NABat transect with forest and urban development covers and landscape index extraction buffers of four spatial scales (B).

### 3.1 Single variable landscape index models

For the single landscape index variable models, total bat activity mostly responded to mixed forest indices positively (Fig. 2). Seven indices describing area, connectivity, and edge had significant effects across all four spatial scales. Four urban indices also had significant effects across all spatial scales, one in each landscape aspect. Total bat activity also responded positively to some deciduous forest indices for area and edge aspects at broad scales. In contrast, evergreen forest indices negatively correlated to total bat activity. Species richness only responded to a few mixed forest indices at broad scales (Fig. 2). Higher species richness was found in areas with higher percentages of continuous mixed forests in less complex shapes with less edge.

**Fig. 2.**
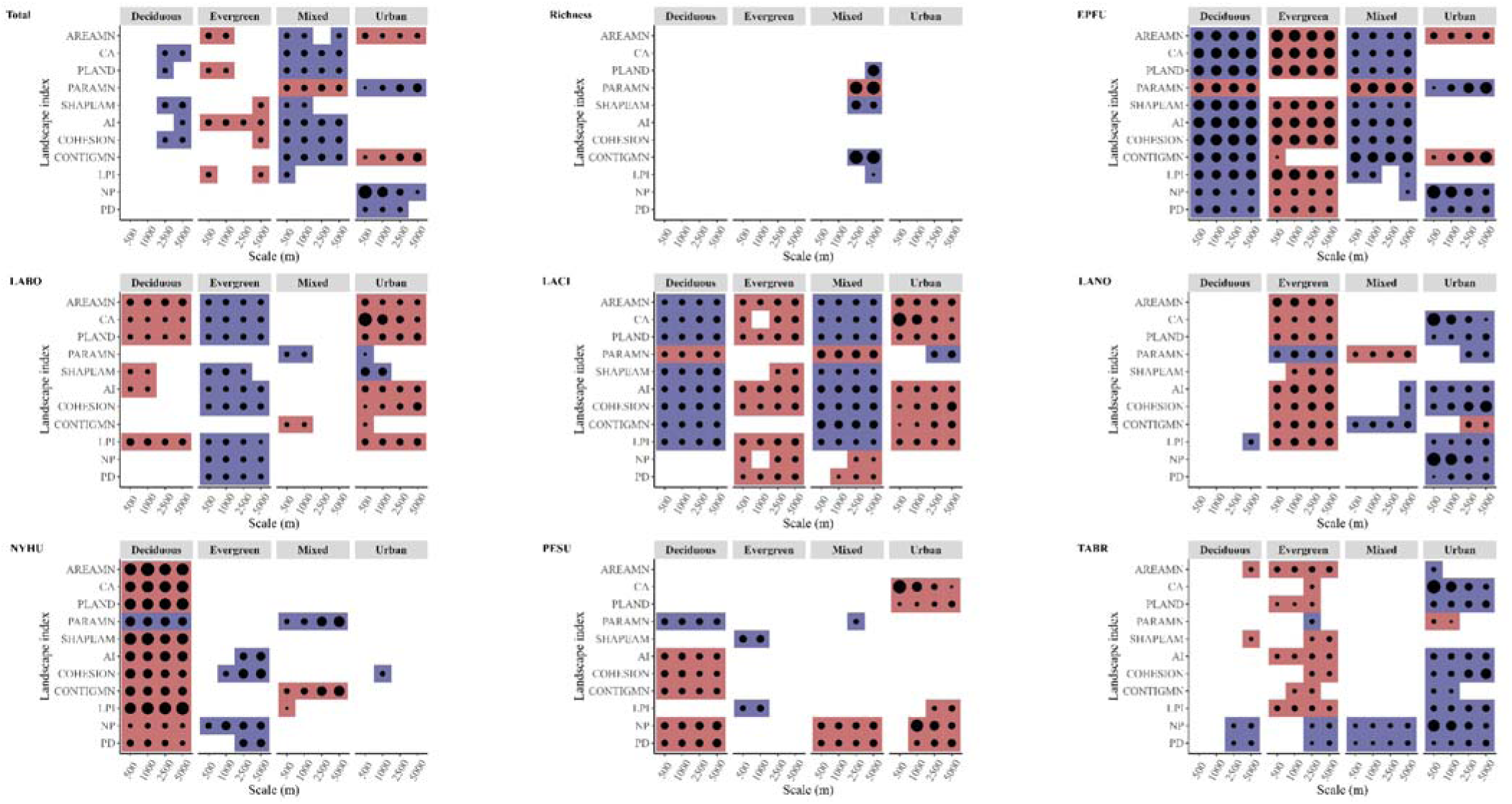
Tile graph for the single variable regression model results of correlation patterns between bat activity and diversity and landscape indices across four spatial scales. A colored tile indicated a significant correlation, blue for positive correlations and red for negative correlations. The dot size represented the absolute value of regression coefficients from models using standardized landscape indices as independent variables.

Species-specifically, big brown bat activity responded to all three types of forests (Fig. 2). Among these, all 11 indices for deciduous forest across all scales correlated to big brown bat activity. Most of these correlations were positive. Similarly, mixed forest indices generally positively correlated with big brown bat activity. In contrast, nine evergreen forest indices across all landscape aspects negatively correlated with big brown bat activity across all spatial scales (Fig. 2). Contradictory to our hypothesis, only a few urban indices correlated with big brown bat activity. Urban indices representing land cover composition did not show correlation with big brown bat activity. However, urban fragmentation indices positively correlated with big brown bat activity. Another known urban affiliated species, the Mexican free-tailed bat, positively responded to seven urban landscape indices across spatial scales (Fig. 2). Mexican free-tailed bat activity negatively responded to several evergreen forest indices, especially those describing connectivity. For the other two types of forests, fragmentation indices were the only noticeable indices correlating to Mexican free-tailed bat activity, exhibiting positive correlations.

The tree roosting silver haired bat showed positive correlations with urban indices for both composition and configuration (Fig. 2). This species also showed negative responses to seven evergreen forest indices. Generally, it did not respond to deciduous forest indices. The other tree roosting species that negatively responded to evergreen forest indices was the hoary bat (Fig. 2). This species also generally negatively responded to urban indices. In contrast, most deciduous and mixed forest indices positively correlated with hoary bat activity, except for fragmentation indices for mixed forest at broad scales. The other three tree roosting species, the eastern red bat, the evening bat, and the tricolored bat generally negatively responded to deciduous forest indices, showing a pattern different from the hoary and silver-haired bats (Fig. 2). The eastern red bat showed most positive responses to evergreen forest indices for both composition and configuration. This species also negatively responded to urban composition and configuration indices, generally consistent with our hypothesis. The evening bat, on the other hand, did not respond to urban indices. Besides deciduous forest indices, this species responded to a few configuration indices for evergreen or mixed forests. The tricolored bat is the only species that did not respond to any forest composition indices (Fig. 2). This species negatively responded to deciduous and mixed forest fragmentation indices across all spatial scales. Urban indices were also negatively correlated with tricolored bat activity.

### 3.2 Multivariate model hierarchical partitioning

The landscape index that contributed the most within a land cover type based on hierarchical partitioning were highlighted in bold in Table 1. Using these landscape indices, we constructed cross-land cover multivariate models. The cross-land cover multivariate model only explained 16% variance in total bat activity (Table 2). Relatively, mean perimeter-area ratio of urban development at the broad scale (5000m) contributed more than all three forest indices combined (Fig. 3, Table 2). This index described the perimeter area ratio of urban edge, showing that total bat activity increased as more urban edge was present, potentially indicating a benefit of urban edge for overall bat abundance (Fig. 4). This was the only instance where an edge index was the top contributing landscape index variable. Due to the limited number of landscape indices that were correlated with species richness, we did not conduct hierarchical partitioning for species richness.

**Fig. 3.**
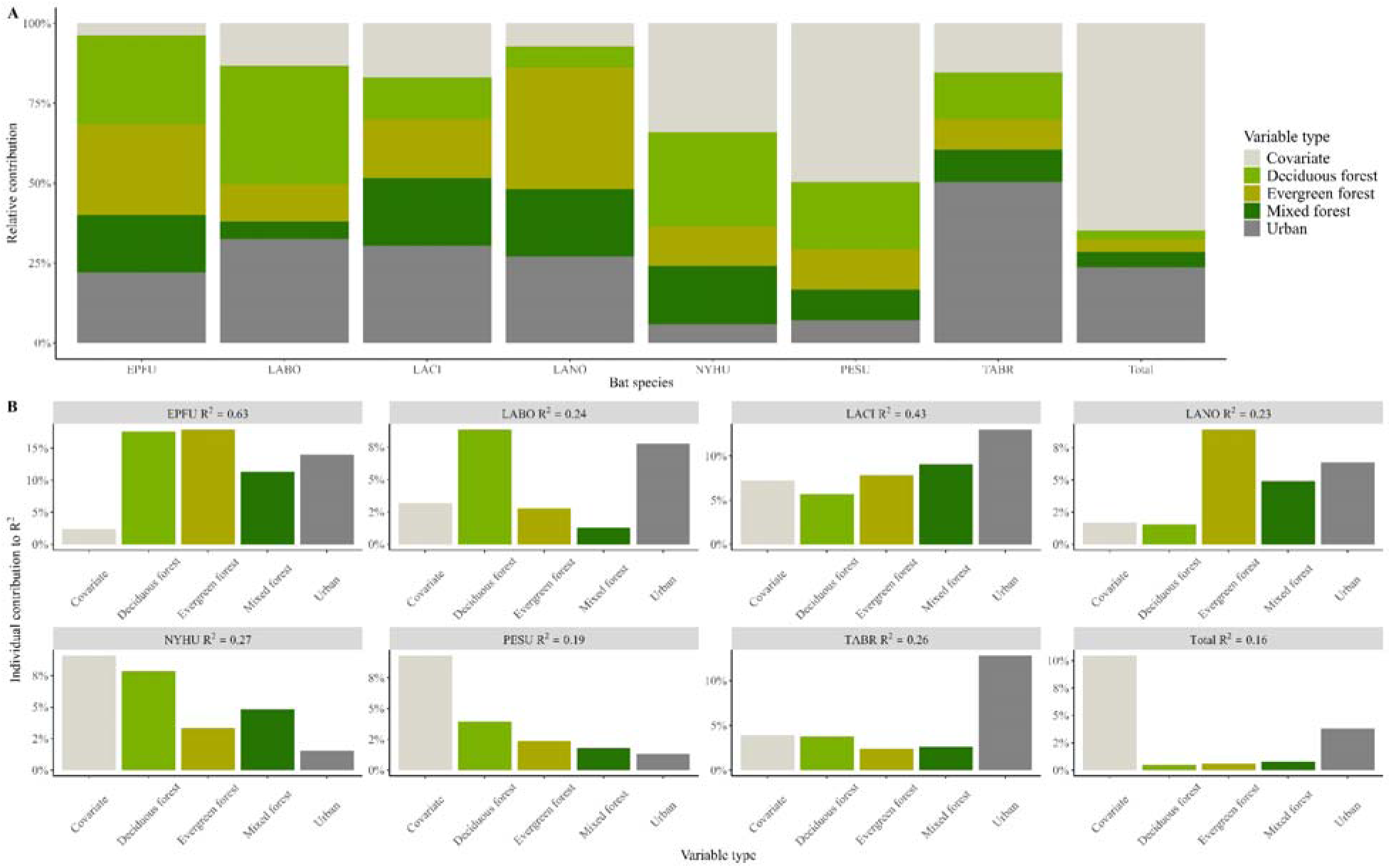
Hierarchical partitioning results on multivariate models of landscape indices across land cover types. (A) relative contribution of individual landscape indices and covariates in predicting bat activity; (B) multivariate model R^2^ (in panel banners) and individual contribution of landscape indices to R^2^.

**Fig. 4.**
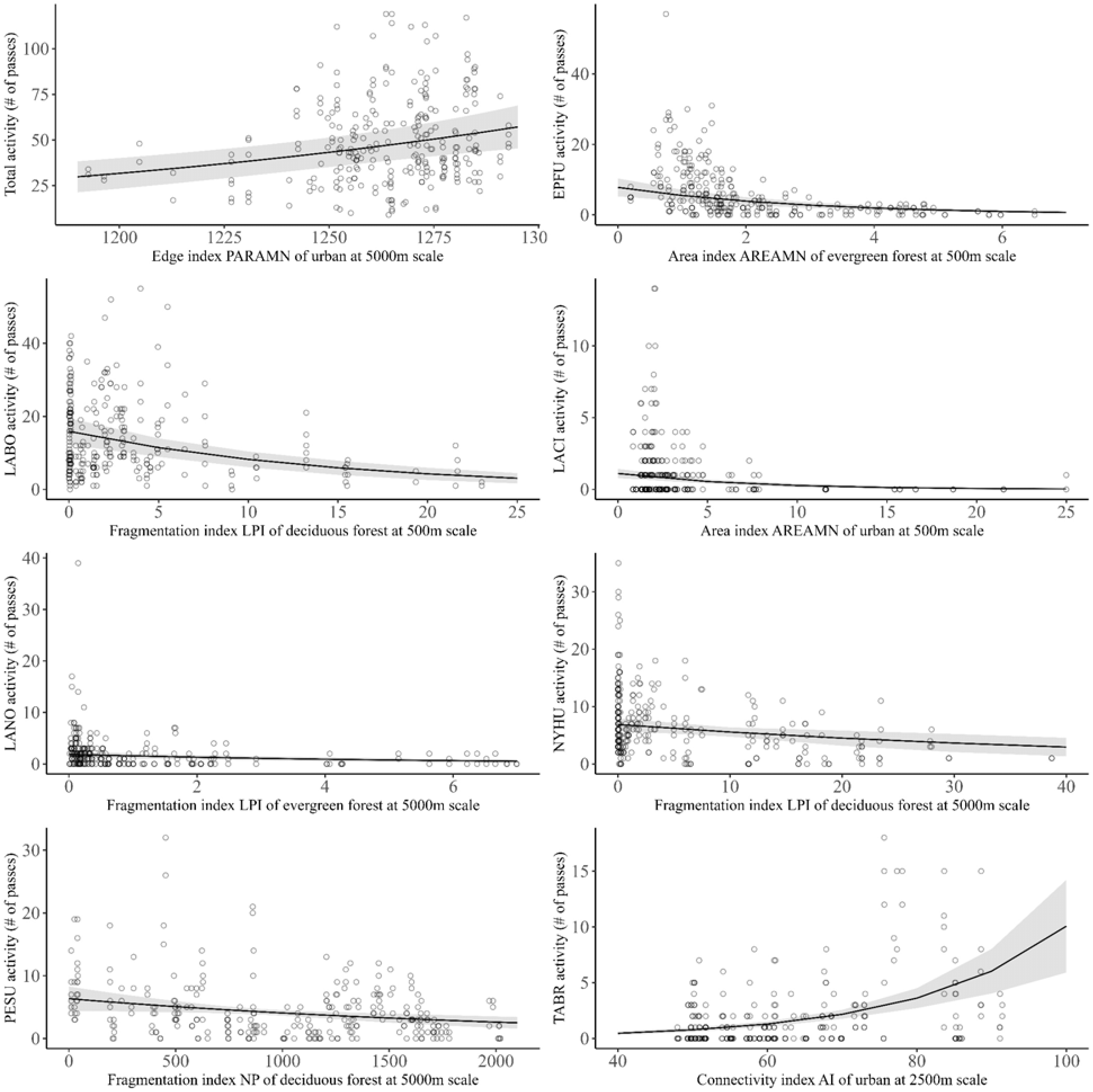
Predicted relationship between the top contributing landscape index (estimated via hierarchical partitioning) and the corresponding bat dependent variable in the cross-land cover multivariate model with raw data points (dots) plotted.

**Table 2.** Cross-land cover multivariate model results, including model R^2^, covariates selected in the base model, and regression coefficients± standard error and the corresponding p value. The top contributing landscape index to R^2^ selected via hierarchical partitioning is highlighted in bold.

| Bat variable | $R^2$ | Covariates | Deciduous | Evergreen | Mixed | Urban |
| --- | --- | --- | --- | --- | --- | --- |
| Total activity | 0.16 | temp + autocov | COHESION-5000<br>-0.048±0.038, 0.211 | AI-500<br>-0.039±0.031, 0.218 | CA-500<br>0.064±0.033, 0.056 | **PARAMN-5000**<br>**0.113±0.033, <0.001** |
| *Eptesicus fuscus* activity | 0.63 | temp | COHESION-1000<br>0.190±0.100, 0.058 | **AREAMN-500**<br>**-0.548±0.090, <0.001** | PARAMN-5000<br>-0.455±0.084, <0.001 | PARAMN-5000<br>0.156±0.081, 0.053 |
| *Lasiurus borealis* activity | 0.24 | autocov | **LPI-500**<br>**-0.338±0.068, <0.001** | SHAPEAM-1000<br>-0.001±0.056, 0.980 | PARAMN-1000<br>-0.036±0.058, 0.534 | PLAND-5000<br>-0.246±0.044, <0.001 |
| *Lasiurus cinereus* activity | 0.43 | temp + wind | LPI-5000<br>0.218±0.137, 0.117 | LPI-5000<br>-0.263±0.134, 0.049 | PARAMN-2500<br>-0.335±0.170, 0.049 | **AREAMN-500**<br>**-0.489±0.185, 0.008** |
| *Lasionycteris noctivagans* activity | 0.23 | temp | LPI-5000<br>0.022±0.116, 0.849 | **LPI-5000**<br>**-0.276±0.094, 0.004** | CONTIGMN-5000<br>0.331±0.133, 0.013 | AI-2500<br>0.384±0.077, <0.001 |
| *Nycticeius humeralis* activity | 0.27 | temp + autocov | **LPI-5000**<br>**-0.192±0.106, 0.070** | PD-5000<br>0.020±0.075, 0.786 | PARAMN-5000<br>0.107±0.074, 0.151 | COHESION-1000<br>0.053±0.052, 0.300 |
| *Perimyotis subflavus* activity | 0.19 | temp + autocov | **NP-5000**<br>**-0.234±0.113, 0.038** | SHAPEAM-1000<br>0.129±0.074, 0.081 | NP-5000<br>0.017±0.117, 0.884 | PD-5000<br>-0.008±0.089, 0.928 |
| *Tadarida brasiliensis* activity | 0.26 | temp + humidity | NP-5000<br>0.483±0.172, 0.005 | CA-2500<br>-0.270±0.106, 0.011 | NP-5000<br>-0.127±0.172, 0.463 | **AI-2500**<br>**0.583±0.089, <0.001** |

For the big brown bat, all four land cover indices each explained more than 10% variance (Fig. 3). Noticeably, evergreen forest mean patch size at the fine scale (500 m) was the most important landscape index variable (Fig. 3, Table 2). Locally, evergreen forest mean patch size negatively correlated with big brown bat activity (Fig.4). Totally, the multivariate model for the big brown bat explained 63% variance of bat activity (Table 2). For the eastern red bat, largest patch index (LPI), describing fragmentation degrees of deciduous forest at the fine scale (500 m) and percentage of urban development, a composition index at the broad scale (5000 m) explained the most of variance in the multivariate model (Table 2). Both indices negatively correlated with eastern red bat activity (largest patch index model result plotted in Fig.4 as it had the highest contribution to R^2^). These results suggested that fewer eastern red bat passes were recorded when an area had a large continuous deciduous forest patch at the fine scale or more urban developments at the broad scale. Urban development mean patch size at the fine scale (500 m), an index for composition, explained more than 10% of variance in hoary bat activity (Fig. 3, Table 2). More large urban development patches along the transects within 500 m would decrease hoary bat activity (Fig. 4).

For the silver-haired bat, evergreen forest configuration index, largest patch index/LPI describing fragmentation, at the broad scale (5000 m) explained more variance than any other forest or urban landscape indices (Fig. 3, Table 2). At the broad scale if an area had a large continuous evergreen forest patch fewer silver-haired bat passes would be recorded (Fig. 4). Similarly, this landscape configuration index also contributed the most in explaining variance in evening bat activity (Fig. 3, Table 2). The only difference was that evening bats responded negatively to large and well-connected deciduous forest patches instead of evergreen forest at the broad scale (5000 m), indicating the species would be more active with fragmented deciduous forests (Fig. 4). In contrast, the tricolored bat might be sensitive to fragmentation in deciduous forest. The fragmentation index, number of patches of deciduous forest at the broad scale (5000 m) explained the most variance in tricolored bat activity (Fig. 3, Table 2) with a negative correlation pattern.

Lastly, aggregation index of urban development at 2500 m scale contributed the most in the multivariate model for Mexican free-tailed bat activity (Fig. 3, Table 2). This was the only instance where a landscape index at the intermediate scale was selected and the only connectivity index as the top contributing variable for R^2^. Aggregation index of urban development at 2500m scale could explain more than 10% of variance in Mexican free-tailed bat activity. When urban development patches at this scale were better connected and aggregated, more Mexican free-tailed bat activity would be recorded (Fig. 4). Collectively for eight cross-land cover multivariate models, two (the big brown bat and the hoary bat) had a composition landscape index as the top contributing variable for R^2^. The rest six models had a configuration landscape index as the top contributing variable for R^2^, four of which were an index describing fragmentation with three being largest patch index/LPI.

## 4. Discussion

The single variable landscape models revealed correlation patterns that were both species- and land-cover-specific, aligning with existing knowledge on how bats respond to varying land covers (Ducci et al. 2015; Mendes et al. 2017; Li et al. 2019). For example, the Mexican free-tailed bat, a known urban affiliated species that commonly depends on anthropogenic structures for roosting (Li and Wilkins 2015; McCracken et al. 2018), predominantly responded positively to urban landscape indices. Bats’ responses to different forest types have been less consistent in the existing literature (Charbonnier et al. 2015; Froidevaux et al. 2021; Węgiel et al. 2023). In the southeastern United States, previous research using all-night acoustic monitoring data indicated that forest types generally had little effect on species-specific activity (Loeb and O’Keefe 2006; Burns et al. 2019). Instead, forest types a more significant role in explaining bat roosting behavior (Drake et al. 2020). Since the fieldwork time window for the NABat mobile transect survey was in the early hours of the night (Martin et al. 2022), our results on bats’ species-specific responses to forest types might partially reflect roosting preferences. For example, foliage-roosting hoary bats in our results showed positive correlations with deciduous or mixed forests and negative correlations with evergreen forest, consistent with their known preference for roosting in deciduous trees (Klug et al. 2012). However, some association patterns in our results were difficult to explain using current knowledge of roost preferences. For example, evening and big brown bats have been shown to exhibit niche overlap in tree roosts in forest ecosystems (Timpone et al. 2006). Yet we found opposite patterns in how these two species responded to deciduous forests. As much of our current understanding of roost selection is based on local-scale studies, our findings highlight the need for broad spatial-scale research on the basic ecology of bats (Pauli et al. 2015).

Spatial scale plays a critical role in shaping how organisms respond to environmental variables (Turner et al. 2001; Fahrig 2017). There are examples of ecological correlation patterns in bats remaining consistent across spatial scales (Li and Kalcounis Rueppell 2018; Li et al. 2019). Regarding how bats respond to varying land covers across spatial scales, existing literature suggests that similar correlative patterns can be found across scales in both urban and forest environments (Ethier and Fahrig 2011; Azam et al. 2016; Mendes et al. 2017; Gallo et al. 2018; Moretto et al. 2019). Similarly, we found consistent correlation patterns across spatial scales, such as how the big brown bat and the evening bat responded to deciduous forest indices or how the red bat responded to evergreen forest indices. Additionally, previous studies have shown that one or two specific spatial scales of land cover tend to have the strongest influence on bat activity or diversity. For example, in Ontario, Canada, Ethier and Fahrig (2011) found that the effect of forest amount on silver-haired bat and eastern red bat activity was strongest at a broad 5 km scale compared to smaller scales. In the Central-North Portuguese coast, the activity of open-space foraging guilds such as *Nyctalus lasiopterus, Eptesicus serotinus,* and *Tadarida teniotis* was most influenced by forest cover at the 1,500 m scale rather than at broader spatial scales (Mendes et al. 2017). Our single variable model results further supported this existing knowledge. For example, the negative effect of urban development on eastern red bat activity was strongest at the 500 m scale. As a tree-roosting species that forages in cluttered spaces (Li and Wilkins 2022), a higher amount of urban development at fine scales may indicate a lack of available vegetated habitat for this species.

An interesting pattern regarding spatial scale and land cover was observed in Brazilian Amazon forests. Rocha et al. (2016) found that frugivorous and gleaning animalivorous bats responded to forest edge and patch densities in divergent ways across scales. While we did not observe similar divergent patterns in our results, several species responded to land cover indices only at broad and fine scales but not at intermediate scales. These results suggested that multiple ecological processes might regulate the observed correlative patterns and highlighted the need for further research to better understand the role of spatial scale in bat ecology (Mendes et al. 2017; Li et al. 2019; López-Baucells et al. 2022).

Our single variable landscape models highlighted that both land cover composition and configuration were important in shaping bat activity and diversity. The effects of urban and forest land covers on bat activity and diversity were well documented in the southeastern United States, reflecting each species’ functional habitat needs (e.g., Loeb and O’Keefe 2006; Hein et al. 2009; Neece et al. 2018; Li et al. 2019). In contrast, very little research has examined the effects of forest or urban development configurations on bat activity and diversity in this region. Brooks et al. (2017) investigated whether forest edge availability affected species-specific bat activity and insect abundance in western North Carolina but found inconclusive results. However, two habitat suitability analyses in the North American temperate forests demonstrated that both forest cover amount and forest edge availability positively correlated with habitat quality for tree-roosting bats (Pauli et al. 2015; Cable et al. 2021). In the temperate-boreal forest transition zone, Ethier and Fahrig (2011) found that forest fragmentation positively affected the abundance of several bat species, including the eastern red bat, a pattern also observed in our study. To our knowledge, our research is the first to present multiple correlation patterns between specific forest/urban development cover configuration indices and bat activity in North American temperate forests. Our results support the consensus that heterogenous landscapes provide better resource complementation for bats; however, key habitats must remain connected to facilitate bat movement among them (Klingbeil and Willig 2009; Hale et al. 2012; Mendes et al. 2017; Shapiro et al. 2020; Falcão et al. 2021; Laforge et al. 2022). For example, we found that the big brown bat generally did not respond to urban development composition indices but showed a positive response to an increased number of urban patches on the landscape. Previous studies have suggested that bats may use urban areas for roosting while foraging outside urban areas or vice versa (Schimpp et al. 2018; Aguiar et al. 2021), indicating the benefits of heterogenous landscapes.

When comparing land cover composition and configuration via multivariate model hierarchical partitioning, we found that for six of the eight bat dependent variables (seven species-specific activity and total bat activity), a land cover configuration index contributed the most to explaining variance. Fragmentation indices of deciduous forest explained the most variance for the eastern red bat, the evening bat, and the tricolored bat, while the evergreen forest fragmentation index explained the most variance for the silver-haired bat. All these species were negatively correlated with forest fragmentation. The impact of forest fragmentation on bats depends on both the species being examined and the surrounding land matrix where fragmentation occurs (Ethier and Fahrig 2011; Rodríguez-San Pedro and Simonetti 2015; Rocha et al. 2016; Brändel et al. 2020). The negative correlation patterns that we identified were consistent with previous research on forest dependent bat species in forest dominated regions. Furthermore, Arroyo-Rodríguez et al. (2016) reported that in the largest Mexican tropical rainforest forest, where intense deforestation has occurred over the past 40 years, forest composition was more important than forest configuration in shaping bat communities. This discrepancy might be related to differences in land matrix and deforestation history between Mexico and our study area. Unlike the Mexican tropical rainforest, our study area has maintained a relatively stable forest cover and has not experienced large-scale deforestation for decades (Arroyo-Rodríguez et al. 2016; Brown and Lambert 2018).

In our research, we examined the forest configurations of three different forest types. For a specific forest type, fragmentation can result from land conversions such as urbanization or agriculture, as well as from silviculture where the introduction of a different forest type disrupts an intact forest. Further research is needed to determine how different causes of fragmentation, particularly silviculture related fragmentation, affect bats at broad scales in forest ecosystems (Loeb 2020). Additionally, while our research examined different forest types that likely had distinct forest structures (Węgiel et al. 2023), we did not include specific forest structure measures in the analysis. Previous studies using detailed forest inventory data demonstrated that factors such as canopy height, vertical complexity, deadwood volume, and other structural variables could play an important role in shaping bat diversity and activity (Novella-Fernandez et al. 2022; Rigo et al. 2024). Future research should explore these factors at broad spatial scales in North America. Beyond forest fragmentation, our hierarchical partitioning model identified urban development connectivity as the most influential factor explaining variance in Mexican free-tailed bat activity. In our land cover data, both urban developments and roads were categorized as urban (Yang et al. 2018). Since no previous research has investigated whether the city layout affected bats, future studies should compare different cities and disentangle the effects of roads from urban developments.

Besides species-specific activity, our research also examined total bat activity and species richness. As several species in our study area have only regional distributions (Grider et al. 2016; Etchison and Weber 2020; Li et al. 2024), we did not expect these collective variables to respond to any specific landscape index. Surprisingly, we found that both the amount and configuration of mixed forest at broad spatial scales correlated with species richness. Additionally, many landscape indices of mixed forest across spatial scales correlated with total bat activity, with the broad spatial-scale urban-nonurban edge index being the most influential predictor, showing a positive correlation with total bat activity. In our study area, oak-pine combinations were the most common tree composition within mixed forests based on previous surveys (Brown and Lambert 2018). These mixed stands can provide diverse roosting opportunities and suitable vertical strata for insect prey preferred by bats, such as beetles and moths (Perry et al. 2007; Maguire et al. 2014; Froidevaux et al. 2021). Similarly, previous studies showed that urban-nonurban edges created heterogenous landscapes that could support high bat abundance by offering diverse resources across different functional habitats (Klingbeil and Willig 2009; Li and White 2024), reinforcing the benefits of habitat heterogeneity and resource complementation discussed earlier. However, given the varying species-specific responses to landscape composition and configuration, we argue that conservation and management decisions must consider species-level information rather than relying solely on collective indicators such as species richness or total bat activity (Klingbeil and Willig 2009; Brändel et al. 2020; Ferreira et al. 2024).

In conclusion, our research demonstrated that both landscape composition and configuration of forests and urban developments influenced bat activity and diversity in a species-specific manner. In North American temperate forest ecosystems, where total forest cover has remained relatively stable over decades, landscape configuration may be more influential than composition and should be considered when designing forest conservation and management plans. We acknowledge that landscape indices, particularly some configuration indices, may not directly translate into on-the-ground conservation and management actions. Therefore, it is essential to evaluate conservation and management strategies (such as forest thinning, road development, and forest restoration) from a landscape perspective, emphasizing how these actions may alter land cover composition and configuration across spatial scales. Additionally, we argue that more basic ecology research (such as roost selection, movement, and diet) is needed for improving our understanding of how bats interact with the environment in both common and rare species. Such knowledge is fundamental to identifying the ecological processes and mechanisms underlying the patterns observed in our research.

## Acknowledgements

The data used in this manuscript were independently collected for various projects with funding support from the United States Fish and Wildlife Service, the North Carolina Wildlife Resources Commission, and the University of North Carolina Greensboro, as part of a collective effort for the North American Bat Monitoring Program (NABat). We sincerely thank all organizations and individuals who have supported and contributed to NABat. The authors’ salaries during this research were supported by their respective institutions, and no direct funding was received for this research. We also thank B. Connor for proofreading early versions of the manuscript.

